# Model of ciprofloxacin subdiffusion in *Pseudomonas aeruginosa* biofilm formed in artificial sputum medium

**DOI:** 10.1101/2020.02.26.966507

**Authors:** Tadeusz Kosztołowicz, Ralf Metzler, Sławomir Wa̧sik, Michał Arabski

## Abstract

We study theoretically and empirically ciprofioxacin antibiotic diffusion through a gel-like artificial sputum medium (ASM) mimicking physiological conditions typical for a cystic fibrosis layer, in which regions occupied by *Pseudomonas aeruginosa* bacteria are present. Our theoretical model is based on the subdiffusion-absorption equation with a fractional time derivative that describes molecules diffusion in a medium structured as Thompson’s plumpudding model; the pudding ‘background’ represents ASM and the plums represent the bacterial biofilm. We show that the process can be divided into three successive stages: (1) only antibiotic subdiffusion with constant biofilm parameters, (2) subdiffusion and absorption of antibiotic molecules with variable biofilm parameters, (3) subdiffusion and absorption in the medium but biofilm parameters are constant. Stage 2 is interpreted as the appearance of an intensive defence bulid-up of bacteria against the action of an antibiotic, in the stage 3 it is likely that the bacteria have been inactivated. Times at which stages change are determined from the experimentally obtained temporal evolution of the amount of substance that has diffused through the ASM with bacteria. Our analysis shows good agreement between experimental and our theoretical results.

## 1 Introduction

Biofilms are defined as complex microbial communities of cells embedded into a matrix of self-produced extracellular polymeric substance (EPS) with increased resistance to antibiotics and host immune response. This is an important issue because in many clinical and industrial settings a biofilm represents a hazardous and costly problem. It is well documented that many chronic infections (65%), particularly those involving medical implants, causing urinary tract infections or in the course of cystic fibrosis involve biofilm-formed bacteria species [4, 10, 13, 14]. Cystic fibrosis (CF) is a disease that causes thick, sticky mucus to build up in the lungs, the digestive tract, and other areas of the body. It is an optimal niche for microorganisms, whose induced chronic lung diseases in children and young adults are evoked mainly by bacteria, such as *Pseudomonas aeruginosa* (80%), *Burkholderia cepacia* and *Staphylococcus aureus*. These bacterial infections lead to progressive pulmonary damage and emphysema. Eradication of bacterial biofilms formed in mucus is a crucial problem because the diffusion of classic antibiotics into biofilm structures is weak and their antibacterial activity might stimulate drug resistance, check Kindler et al. [5] for additional references. Biophysical properties of biofilm structure EPS in gel-like mucus are directly associated with reduced susceptibility to antibiotics and limit the effective eradication of bacteria [16].

In this study, we focus on an experimental system mimicking physiological conditions typical to cystic fibrosis to measure antibiotic (ciprofioxacin) transport through a *P. aeruginosa* biofilm. Ciprofioxacin antibiotic is one of the most commonly used fiuoroquinolone in the treatment of *P. aeruginosa* infections during cystic fibrosis. We present a model of the antibiotic diffusion in a biofilm that can be described as a plumpudding. Because the matrix (the pudding representing the cystic fibrosis EPS) has a gel-like consistency and has physicochemical interactions with the diffusing antibiotic, subdiffusion may occur in this medium. In the experimental system pudding represents an artificial sputum medium (ASM). The plums correspond to bacterial cells making up microcolonies, such as *Pseudomonas aeruginosa.* The antibiotic only acts on bacteria located in the plums. As a result of the activation of bacterial defence mechanisms, an antibiotic particle can be permanently or temporarily retained in the plums or can be destroyed, depending on the type of bacterial defence mechanism. The process that a molecule can be permanently trapped or destroyed is effectively described by the subdiffusion-absorption equation. The process of temporary retention of a molecule can be regarded as a reversible reaction. In this case, the process is described by the subdiffusion equation in which the reaction term is absent.

One of the main problems in the treatment of bacterial diseases is to check whether an antibiotic with a given initial concentration affects the bacteria, and if so, to determine the time after which the bacteria stop defending themselves against the effects of an antibiotic. It can be assumed that then the bacteria have been inactivated. In order to find the answer to this problem we study experimentally the time evolution of the amount of antibiotic that has diffused through the biofilm *W*_*B*_ in a system presented schematically in Fig. 1. The theoretical description is derived from a model based on the subdiffusion-absorption equation with fractional time derivative which describes the process in the biofilm [7]. The theoretical model shows that the form of *W*_*B*_ depends on the presence of antibiotic molecule absorption in the biofilm. The appearance of absorption of antibiotic molecules and/or a change in biofilm parameters indicates that the bacteria sense the antibiotic and activate their defence mechanism against the antibiotic. Absorption of antibiotic in the biofilm is related to the intensity of antibiotic-bacterial interactions. Analysis of the function *W*_*B*_ obtained experimentally allows us to estimate the time after which the bacteria will react to the antibiotic action and the time when the bacteria cease to react.

**Figure 1:**
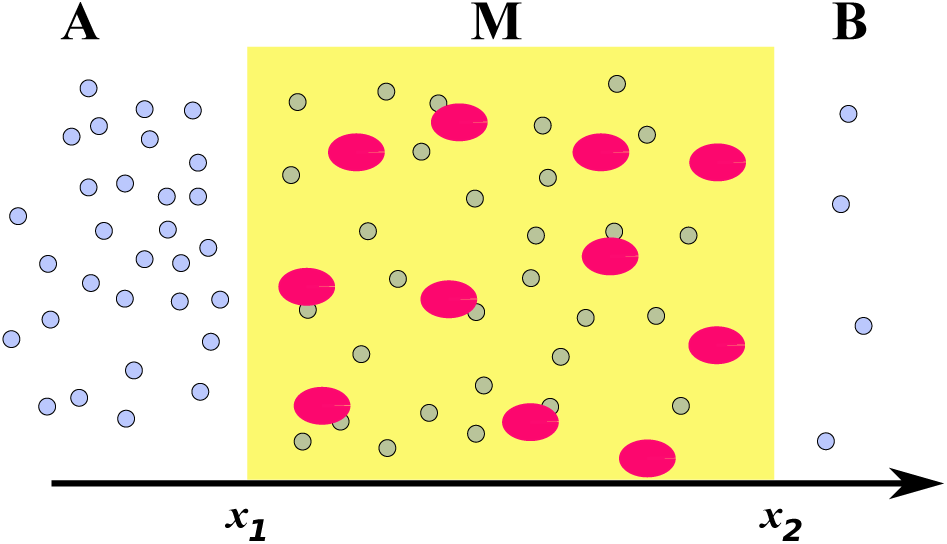
Diffusion of antibiotic molecules through a biofilm that looks like a plumpudding. The system consists of diffusive media *A* and *B* and subdiffusive biofilm *M*. The dark regions in *M* (plums) represent bacterial cells forming microcolonies which can absorb the antibiotic molecules. The pudding (yellow) represents the cystic fibrosis biofilm.

## 2 Materials and nethods

### 2.1 Theory

We consider antibiotic diffusion in the model system shown in Fig. 1. We assume that the system is homogeneous at a plane perpendicular to the *x* axis, thus the problem is one-dimensional. The system consists of the three regions *A* (*x < x*_1_), *M* (*x*_1_ *< x < x*_2_), and *B* (*x > x*_2_); these symbols also label the concentrations in the relevant parts of the system. At the initial moment part *A* contains a homogeneous aqueous solution of antibiotic (ciprofioxacin), while part *B* is filled with pure water. The equations which describe the process are the normal diffusion equation in regions *A* and *B*

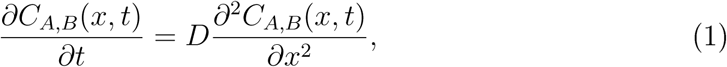

and the time–fractional diffusion–reaction equation

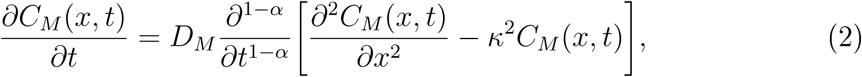

where *α, D*_*M*_, and *κ* denote the subdiffusion parameter, the subdiffusion coefficient, and the absorption coefficient, respectively, in the medium *M*. *D* is the normal diffusion coefficient in media *A* and *B*. Eq. (2) contains the Riemann–Liouville fractional time derivative of order 1 − *α* [12]. Using Eq. (2) to describe diffusion-absorption in a biofilm, we use the approximation of a homogeneous medium *M*. The parameters *α, D*_*M*_, and *κ* are effective parameters describing diffusion and absorption throughout the medium; they represent the antibiotic diffusion through the matrix and they interaction with the plums. The initial concentration is *C*_*A*_(*x*, 0) = *C*_0_, *C*_*M,B*_(*x*, 0) = 0. The boundary conditions at the border between the media, which were derived in [7], are 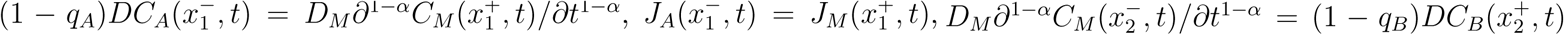, and 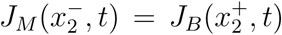, where the diffusive fiuxes are defined as *J*_*A,B*_(*x, t*) = −*D∂C*_*A,B*_(*x, t*)*/∂x* and *J*_*M*_ (*x, t*) = −*D*_*M*_ (*∂*^1−*α*^*/∂t*^1−*α*^)*∂C*_*M*_ (*x, t*)*/∂x*. The boundary conditions mean that the diffusion fiux at the biofilm surface is continuous and that the antibiotic molecule that tries to leave the biofilm can do so without any obstacles, but its passage into the biofilm surface in the opposite direction can be made with probabilities 1 −*q*_*A*_ (for the biofilm surface located at *x*_1_) and 1−*q*_*B*_ (at *x*_2_) [7, 8]. Since the concentration of the antibiotic is higher at the *x*_1_ surface and changes faster than at the *x*_2_ surface, activated bacterial defence mechanisms impede diffusion of the antibiotic through the *x*_1_ surface, *q*_*A*_ *> q*_*B*_.

In the experiment, we determine the temporal evolution of the function 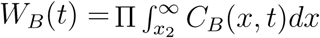, where Π is the biofilm surface area. In [7] it was shown that the form of the function *W*_*B*_ depends on whether or not absorption takes place in the medium *M*. Four stages of the diffusion process of the antibiotic through the biofilm were distinguished, taking into account the following ‘physical’ criteria: (1) whether or not absorption of antibiotic molecules in the biofilm is observed, and (2) if the biofilm parameters are constant or if at least one of them changes over time. Since the function *W*_*B*_ is qualitatively different at each stage, it serves to identify these stages in the process under consideration. For a sufficiently long time, this function takes the following forms when the biofilm parameters are constant

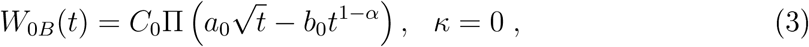

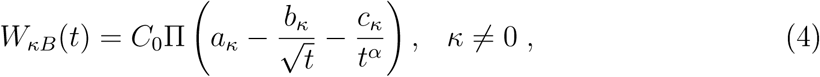

where 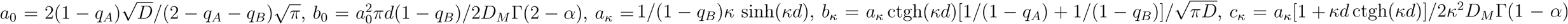, and *d* = *x*_2_ − *x*_1_. Vhen absorption occurs and the biofilm parameters change over time, the following function has been proposed [7] 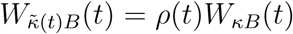, where *ρ*(*t*) is to be determined from empirical data. For the case considered in this paper, the function *ρ*(*t*) = *a*−*b/t* gives a good agreement of the *W*_*B*_ function and experimental results in the time interval ⟨*t*_1_, *t*_2_⟩. The times *t*_1_ and *t*_2_ separating the subsequent stages are defined by the equations *W*_0*B*_(*t*_1_) = _*κB*_(*t*_1_) and 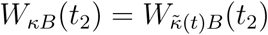. Thus, when *κ* ≠ *const.* ≠ 0, *W*_*B*_ takes the form

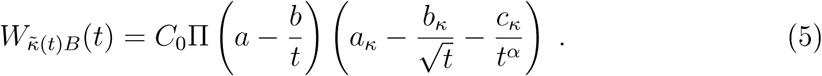

The parameters *a*_*κ*_, *b*_*κ*_, and *c*_*κ*_ for the functions Eqs. (4) and (5) are the same. The stage without absorption and changing biofilm parameters has not been observed here.

The parameter *α* in Eqs. (3), (4), and (5) is the same, so we assume that this parameter does not change over time. We suppose that the probability of absorption of antibiotic molecules per unit length of biofilm, which is controlled by the parameter *κ*, and the biofilm thickness are small, *κd* ≪ 1 [7]. Assuming that the parameters *D* and *α* are constant, by swapping the other parameters in Eq. (4) according to the formula

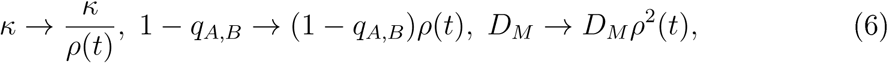

we obtain Eq. (5).

### 2.2 Experiment

We determine the temporal evolution of the function *W*_*B*_(*t*). This function was calculated numerically from experimental concentration profiles *C*_*B*_. The measurements of the concentration profiles were conducted in a vertically oriented vessel consisting of two glass cuboidlike cuvettes (70 mm high, 10 mm wide, 65 mm long) separated by a hori:lontally located PET membrane with PAO1 biofilm. The *P. aeruginosa* PAO1 biofilm was formed for 96 h at 37° C in artificial sputum medium (ASM) on the PET membrane with pore diameter 1 *µ*m, as an element of BD Falcon™ Cell Culture Inserts. ASM was formulated to mimic the sputum of cystic fibrosis (CF) patients and to study microbial biofilms, for example, *Pseudornonas aeruginosa* coloni:lation in CF lungs [6]. At the first step of the analysis, the percentage of the membrane covered by PAO1 biofilm was estimated. The images of the membrane covered by PAO1 biofilm were stained by CV (0.004%) for 15 min, converted to grey-scale digital images and analy:led with the ImageJ computer imaging software program [15]. Initially, the upper cuvette was filled with an aqueous solution of ciprofioxacin with *C*_0_ = 1 mg*/*ml = 3.02 mol*/*m^3^, whereas the lower cuvette was filled with pure water. Since the concentration gradient is solely in the vertical direction, antibiotic transport is effectively one dimensional. The substance concentration was measured by means of laser interferometry, the method and the setup used in the experiment are described in more detail in [2, 19]. Due to technical reasons, the measurement of the antibiotic concentration can be conducted only in the cuvette region *B*.

## 3 Results

In Fig. 2 we present the amount of substance *W*_*B*_ that fiows into part *B*. Points represent the experimental data, which have been calculated from the experimentally measured antibiotic concentration profiles, and lines represents the theoretical functions Eqs. (3)–(5). We fit the theoretical functions to the experimental data and find the values of the parameters given in the figure caption. In all cases, the subdiffusion coefficient *α* = 0.96 is the same. This parameter has been determined as one of the fitting parameters when analy:ling Stage 1. Unfortunately, the statistic is too poor to determine the error for *α*. Because the consistency of ASM is very similar to the consistency of 1% aqueous agarose solution for which *α* = 0.95 [9], we suppose that there is subdiffusion in the ASM medium and the value of the parameter *α* is realistic.

**Figure 2:**
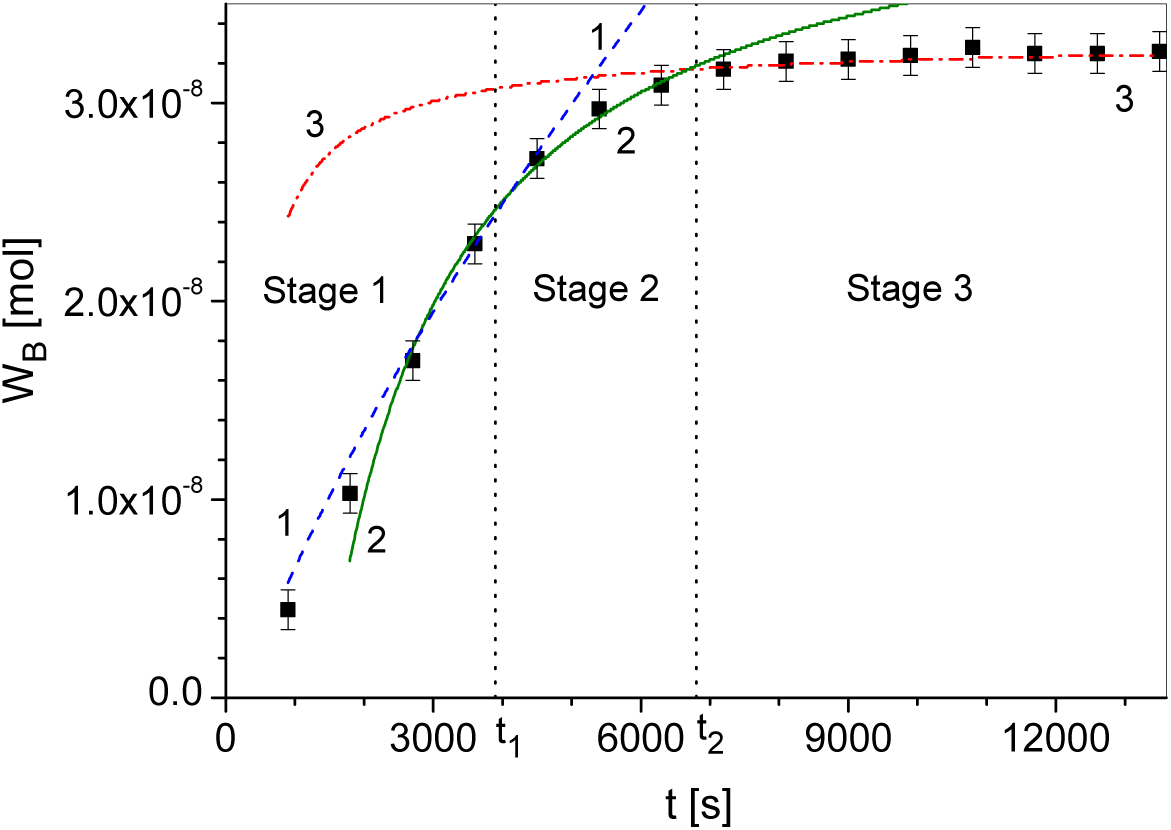
The amount of ciprofioxacin which diffuses into part *B*. Points represent the experimental data and lines represent theoretical functions. Line No. 1 (dashed line) corresponds to function *W*_0*B*_ Eq. (3) for 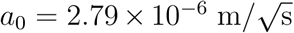 and *b*_0_ = 4.21 *×* 10^−5^ m*/*s^0.04^, line No.2 (solid line) represents 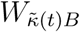 Eq. (5) for *a*_*κ*_ = 1.59 × 10^−4^ m, 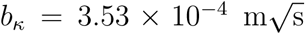 and *c*_*κ*_ = 2.22 × 10^−2^ m s^0.96^, *a* = 1.28, and *b* = 1850 s, line No. 3 (dashed-dotted line) represents *W*_*κB*_ Eq. (4), *t*_1_ = 3900 s, *t*_2_ = 6800 s, *C*_0_ = 3.02 mol*/*m^3^, and Π = 7.0 *×* 10^−5^ m^2^. For all cases *α* = 0.96. The two vertical dotted lines are the boundaries between the stages.

The observed order of stages in Fig. 2 is as follows: initially we see *Stage 1* described by Eq. (3), followed by *Stage 2* described by Eq. (5), and finally we see *Stage 3* described by Eq. (4). We note that the function *ρ*(*t*) is increasing over time. Due to Eq. (6), the parameter *κ* is a function decreasing over time and the parameter *D*_*M*_ increases over time in Stage 2. Changes in parameter values are: *κ* decreased by 19% and *D*_*M*_ increased by 51% in the time interval ⟨*t*_1_, *t*_2_⟩. The interpretation of the process is that at Stage 1 there is a sub-inhibitory concentration of the antibiotic near the bacteria located in the plums. The bacterial defence mechanisms are then not fully activated. In Stage 2 the antibiotic concentration near the plums increases, the defence mechanisms is activated. The antibiotic effect weakens the bacteria to such a level that the infiuence of factors causing absorption of antibiotic molecules and slowing down of diffusion decreases, which causes following changes in the biofilm parameters. At the moment *t*_2_ these parameters stop changing, which can be interpreted as the inability of bacteria to continue active defence.

## 4 Discussion

A different course of the process has been observed when a ‘dense biofilm’ attacked by bacteria fills the entire space of the central part *M* of the system. In [7] it is shown that in this case the order of the stages is as follows (we keep the notation used in the present paper): Stage 1, Stage 3, then Stage 2. The function *ρ*(*t*) is decreasing, so the absorption coefficient increases while the subdiffusion coefficient decreases over time. The reason that the course of antibiotic diffusion in a dense biofim and in the plumpudding scenario is qualitatively different is as follows. In a dense biofilm, antibiotic molecules can feel the action of the bacterial defence mechanisms the whole time inside the biofilm. In a plumpudding case, an antibiotic molecule feels the defence mechanism after reaching a plum occupied by bacteria. This plum can be attacked intensively, from all sides, by an antibiotic which concentration at the surface may increase over time due to diffusion process. This leads to a weakening of the bacterial defences faster than in a dense biofilm. Since there are a lot of mechanisms for the defence of bacteria against the effects of an antibiotic [1], whether the order of stages in both cases is universal is the open question.

In summary, we have shown a method to experimentally check whether a biofilm in a plumpudding scenario absorbs antibiotic particles and whether the biofilm parameters change over time. The presented three-stage model describes subdiffusion of ciprofioxacin through ASM with *Pseudomonas aeruginosa* bacteria. However, studying other processes it is possible to attach the fourth stage in which there is no absorption and the parameters of the biofilm change over time; this stage is not observed in Fig. 2. The division into stages is based on the physical aspects of the process. It is not obvious which bacterial defence mechanisms can be included in these stages, see [1, 3, 11, 17, 18]. However, we believe that determining the order of the stages may facilitate the biological interpretation of the process for various antibiotic-biofilm systems. The presented method may be useful when planning treatment for cystic fibrosis patients who also have another bacterial infection. In particular, this method allows to determine the time at which the bacteria intensify the process of defence against the antibiotic and the time at which this process is stopped.

## Acknowledgenents

This paper was partially supported by the Polish National Science Centre under grant no. UMO-2016/21/B/NZ6/01157.

## References

[1] Anderson GG, O’Toole GA. 2008. Innate and induced resistance mechanisms of bacterial biofilms. Current Topics in Microbiology and Immunology 322:85–105. DOI: 10.1007/978-3-540-75418-3 5, PMID: 18453273.

[2] Arabski M, Wąsik S, Dworecki K, Kaca W. 2007. Laser interferometric determination of ampicillin and colistin transfer through cellulose biomembrane in the presence of Proteus vulgaris O25 lipopolysaccharide. Journal of Membrane Sciences 299:268–275. DOI:10.1016/j.memsci.2007.05.003.

[3] Chambless JD, Hunt SM, Stewart PS. 2006. A three-dimensional computer model of four hypothetical mechanisms protecting biofilms from antimicrobials. 2006. Applied and Environmental Microbiology 72:2005–2013. DOI: 10.1128/AEM.72.3.2005-2013.2006, PMID: 16517649.

[4] Costerton JW. 2001. Cystic fibrosis pathogenesis and the role of biofilms in persistent infection. Trends in Microbiology 9:50–52. DOI: 10.1016/s0966-842x(00)01918-1, PMID: 11173226.

[5] Kindler O, Pulkkinen O, Cherstvy AG, Metller R. 2019. Burst statistics in an early biofilm quorum sensing model: the role of spatial colony-growth heterogeneity. Scientific Reports 9:12077 (19p). DOI: 10.1038/s41598-019-48525-2, PMID: 31427659.

[6] Kirchner S, Fothergill JL, Vright EA, James CE, Mowat E, Vinstanley C. 2012. Use of artificial sputum medium to test antibiotic efficacy against Pseudomonas aeruginosa in conditions more relevant to the cystic fibrosis lung. Journal of Visualized Experiments 64:e3857 (2012). DOI: 10.3791/3857, PMID: 22711026.

[7] Kosztolowicz T, Metller R. 2020. Diffusion of antibiotics through a biofilm in the presence of diffusion and absorption barriers arXiv: 2003.01516 [physics.bio-phys].

[8] Kosztolowicz T. 2019. Model of anomalous diffusion-absorption process in a system consisting of two different media separated by a thin membrane. Physical Review E 99:022127 (16p). DOI: 10.1103/PhysRevE.99.022127, PMID: 30934262.

[9] Kosztolowicz T. 2017. Subdiffusion in a system consisting of two different media separated by a thin membrane. International Journal of Heat and Mass Transfer 111:1322–1333. http://dx.doi.org/10.1016/j.ijheatmasstransfer.2017.04.058.

[10] Lesouhaitier O, et al. 2019. Host peptidic hormones affecting bacterial biofilm formation and virulence. Journal of Innate Immunity textbf 11:227–241. DOI: 10.1159/000493926, PMID: 30396172.

[11] Mah TFC, O’Toole GA. 2001. Mechanisms of biofilm resistance to antimicrobial agents. Trends in Microbiology 9:34–39. DOI: 10.1016/s0966-842x(00)01913-2, PMID: 11166241.

[12] Metzler R, Klafter J. 2000. The random walk’s guide to anomalous diffusion: a fractional dynamics approach. Physics Reports 339:1–77. DOI: 10.1016/s0370-15730000070-3.

[13] Parker MR, Tolker-Nielsen T. 2008. Pattern formation in Pseudomonas aeruginosa biofilms. Current Opinion in Microbiology 11:560–566. DOI: 10.1016/j.mib.2008.09.015, PMID: 18935979.

[14] Sauer K, Camper AK, Ehrlich GD, Costerton JW, Davies DG. 2002. Pseudomonas aeruginosa displays multiple phenotypes during development as a biofilm. Journal of Bacteriology 184:1140–1154. DOI: 10.1128/jb.184.4.1140-1154.2002, PMID: 11807075.

[15] Schneider CA, Rasband WS, Eliceiri KW. 2012. NIH Image to ImageJ: 25 years of image analysis. Nature Methods 9:671675. DOI: 10.1038/nmeth.2089, PMID: 22930834.

[16] Stewart PS. 2003. Diffusion in biofilms. Journal of Bacteriology 185:1485–1491. DOI: 10.1128/jb.185.5.1485-1491.2003, PMID: 12591863.

[17] Stewart PS. 1994. Biofilm accumulation model that predicts antibiotic resistance of Pseudomonas aeruginosa biofilms. Antirnicrobial Agents and Chernotherapy. 38:1052–1058. DOI: 10.1128/aac.38.5.1052. PMID: 8913456.

[18] Stewart PS. 1996. Theoretical aspects of antibiotic diffusion into microbial biofilms. Antirnicrobial Agents and Chernotherapy. 40:2517-2522. PMID: 8913456.

[19] Wąsik S, Arabski M, Drulis-Kawa Z, Gubernator J. 2013. Laser interferometry analysis of ciprofioxacin and ampicillin diffusion from liposomal solutions to water phase. European Biophysics Journal 42, 549–558. DOI: 10.1007/s00249-013-0904-2.

